# A stable mode of bookmarking by TBP recruits RNA Polymerase II to mitotic chromosomes

**DOI:** 10.1101/257451

**Authors:** Sheila S. Teves, Luye An, Aarohi Bhargava-Shah, Liangqi Xie, Xavier Darzacq, Robert Tjian

## Abstract

**How a cell maintains transcriptional fidelity across mitosis has remained an enduring mystery in biology. One challenge arises during mitosis when chromatin becomes condensed and transcription is shut off. How do the daughter cells re-establish the original transcription program? Here, we report that the TATA-binding protein (TBP), a key component of the core transcriptional machinery, remains bound globally to active promoters in ES cells during mitosis. Using live-cell single-molecule imaging, we observed that TBP mitotic binding is highly stable, with an average residence time of minutes. This stable binding is in stark contrast to typical TFs with residence times of seconds. To test the functional effect of mitotic TBP binding, we used a drug-inducible degron system and found that TBP promotes the association of RNA Polymerase II with mitotic chromosomes, and facilitates transcriptional reactivation following mitosis. These results suggest that the core transcriptional machinery maintains global transcriptional memory during mitosis.**

## INTRODUCTION

Faithful maintenance of transcription programs is essential to direct cell state and identity. However, during each cell cycle event of mammalian cells, this transcriptional fidelity is challenged in multiple ways. First, the genome becomes highly condensed into characteristic mitotic chromosomes (1). Secondly, transcription becomes globally inhibited through cell cycle dependent phosphorylation cascades that inactivate the transcriptional machinery (2–4). Lastly, the disassembly of the nuclear membrane combined with TF exclusion from mitotic chromosomes was thought to result in the dispersal of most TFs throughout the cytoplasm (5–8). How then do the new daughter cells faithfully re-establish the original transcription program? A clue to the potential mechanism for transcriptional memory through the cell cycle emerged when the first TF that remained bound to mitotic chromosomes was identified (9), and a hypothesis for mitotic bookmarking was proposed (10). The theory posited that a select group of privileged TFs maintain the ability to bind to target sequences during mitosis and “bookmark” their target genes for efficient reactivation following cell division. Recently, we discovered that previous evidence for the widely observed exclusion of TFs from mitotic chromosomes was an artifact of chemical crosslinking-based cell fixation prior to imaging, e.g., formaldehyde crosslinking (11). By relying on live cell imaging, our studies indicated that in fact most TFs remain associated with mitotic chromosomes but in a dynamic manner. Rather than acting as a stable bookmark, sequence-specific TFs seem to ‘hover’ on mitotic chromosomes in general. However, it was not clear how such a dynamic mode of bookmarking by TFs could maintain stable transcriptional memory during mitosis.

In order to effect transcriptional regulation, sequence-specific TFs at enhancers must cross-talk with the RNA Polymerase II (Pol II) machinery at promoters. Indeed, previous studies have shown that promoters and transcription start sites (TSSs) of certain genes remain sensitive to permanganate oxidation during mitosis (10), providing indirect evidence for potential promoter-specific bookmarking factors that maintain an open chromatin conformation at these chromosomal locations. Central to the recruitment of Pol II to promoters is the TATA-binding protein (TBP), the prototypic member of the multi-subunit core promoter recognition ensemble TFIID (12). At most eukaryotic promoters, TBP and its associated factors (TAFs) are the first components of the pre-initiation complex (PIC) to bind promoters of active genes in a stepwise assembly of the general TFs and Pol II. An essential protein, TBP forms extensive contacts with promoter DNA by introducing sharp kinks that results in an almost 90° bend in the double helix, which leads to partial DNA unwinding and subsequent widening of the minor groove for maximum TBP contact (13–15). TBP also plays a crucial role in the hierarchical assembly of the transcriptional machinery (16), and in the positioning of promoter DNA into the Pol II binding site (17, 18). These essential functions at the promoter make TBP an attractive candidate for a global facilitator of transcriptional memory. In order for TBP to be considered a global bookmarker of active genes, TBP must (1) bind to promoters of active genes during mitosis, and (2) promote efficient transcriptional reactivation following mitosis.

Over time, a number of studies have tested the hypothesis of TBP as a mitotic bookmarker, but results have been conflicting. For instance, immunofluorescence (IF) staining of TFIID subunits, including TBP, showed cellular redistribution of the subunits away from chromosomes during mitosis (19), while live-cell imaging of GFP-tagged but over-expressed TBP shows enrichment on chromosomes throughout mitosis (20). We now know, however, that these conclusions need to be revisited, as these early studies suffered from previously unrecognized experimental artifacts, including the formaldehyde crosslinking artifact we recently uncovered (11), as well as over-expression biases for proteins with highly regulated concentrations within the cell (21). Furthermore, some studies have found that TBP remains bound at specific loci during mitosis via chromatin immunoprecipitation (22), whereas others have not (23). These conflicting results suggest that the question of whether TBP bookmarks active gene promoters remains unresolved. Moreover, the functional consequence of TBP bookmarking (i.e. promoting transcriptional memory through mitosis), if any, has thus far not been established.

Through live-cell imaging of endogenously tagged proteins, we report that TBP stably binds to chromatin for minutes in contrast to most sequence-specific TFs studied thus far that bind on the order of seconds. Moreover, TBP remains stably associated with mitotic chromosomes, and such stable binding is also in contrast to the more dynamic mode of studied sequence-specific TF association with mitotic chromosomes (11). TBP stable binding occurs on a global scale at promoters of active genes during mitosis. We dissected the functional consequence of TBP binding during mitosis by using a drug inducible protein degradation system and found that TBP recruits a small fraction of Pol II molecules to mitotic chromosomes. Lastly, nascent RNA analysis indicates that TBP promotes efficient reactivation of global ES cell transcription program following mitosis. TBP, therefore, is unique among the TFs we have studied thus far, as it appears to stably bookmark active genes during mitosis and thereby maintain transcriptional memory across the cell cycle.

## RESULTS

### TBP bookmarks mitotic chromosomes of mouse ES cells

A previous study has shown that DNase I hypersensitive sites at promoters are maintained during mitosis in murine erythroblast cells (24). Similarly, we have found that mitotic chromosomes of mouse ES cells remain as accessible to Tn5 transposase insertion as interphase chromatin (11). To examine chromatin accessibility specifically at the TSS of mouse ES cells, we analyzed our previously published ATAC-seq data at the TSS of all genes. Reads under 100 bp indicate potential transcription factor (TF) binding sites, and these sites are highly enriched at the TSS of all active genes in asynchronous cells (Figure 1A). Notably, these reads remain nearly equally enriched at the TSS of active genes during mitosis (Figure 1A) and the level of enrichment remains proportional to expression levels as seen by heatmap analysis. When we averaged the signal across all genes, we observed that the ATAC-seq signal at the TSS in mitosis is about 91% of the signal in asynchronous cells (Figure 1A, bottom). Furthermore, when we performed unbiased k-means clustering, we do not observe any population differing between the two samples (Figure 1 - Figure Supplement 1), suggesting high concordance in accessibility at the TSS between asynchronous and mitotic cells. Enrichment of short fragments by ATAC-seq at a given locus is indicative of TF binding. Such maintenance of accessibility at the TSS suggests that the TSS of active genes may be bookmarked by TFs throughout mitosis. We also analyzed the mono-nucleosome sized fragments (180 — 250 bp) from the ATAC-seq at the TSS genome-wide. Remarkably, we found that the accessibility of the nucleosomes flanking the TSS increased during mitosis relative to interphase chromatin (Figure 1B). When we averaged the signal for all genes, we observed a 15% increase in signal in mitotic cells (Figure 1B, bottom). Furthermore, unbiased k-means clustering showed that, while some variation between clusters exist, mitotic signal is consistently higher than the asynchronous signal (Figure 1 - Figure Supplement 2). These results suggest that nucleosomes around the TSS may have a more open and accessible configuration during mitosis than in interphase, despite the general inhibition of transcription observed in mitosis (2).

**Figure 1.**
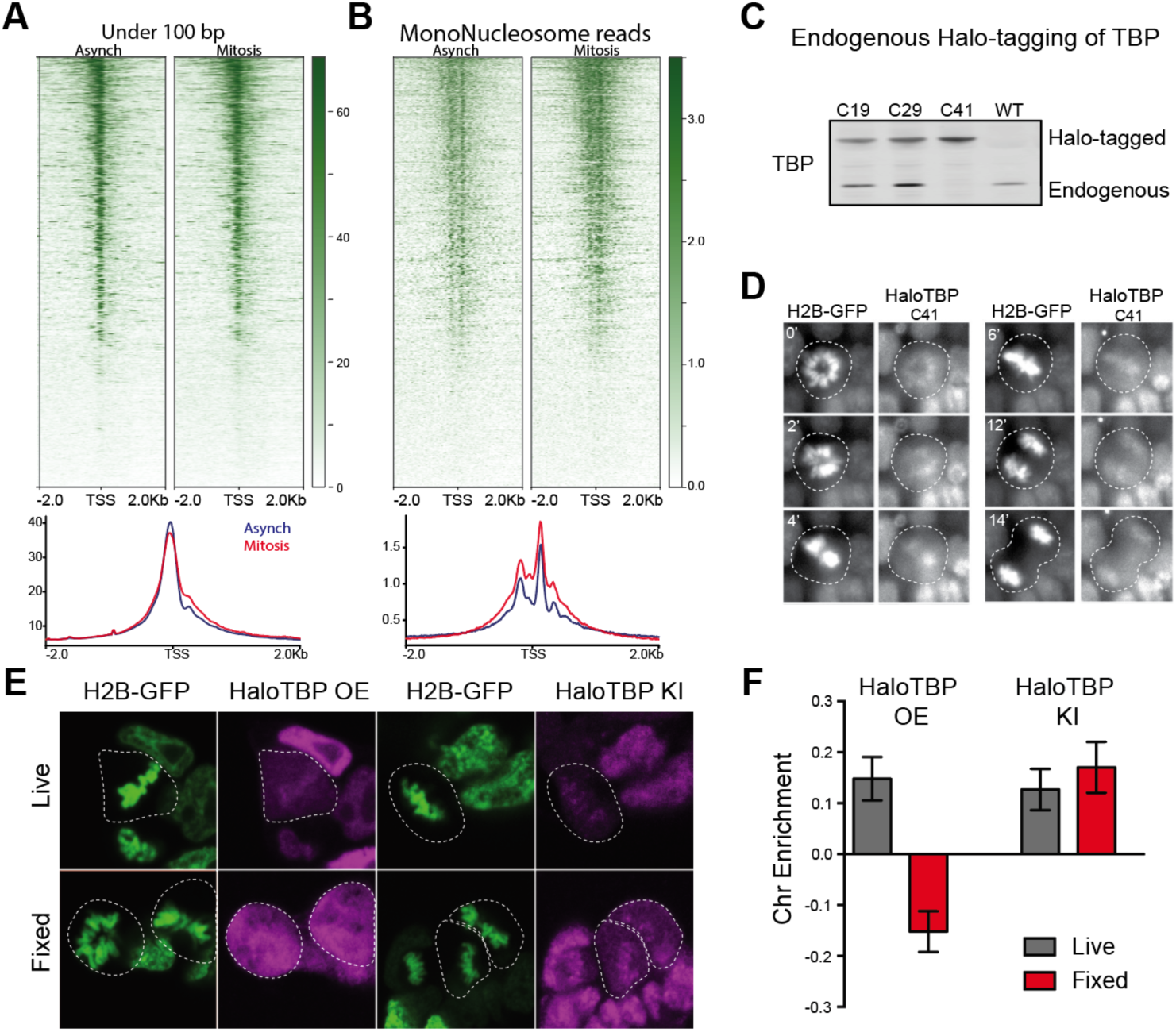
Endogenous TBP associates with mitotic chromosomes. **A**. Previous ATAC-seq data (11) examined for TSS analysis. Heatmaps for all genes centered at the TSS (top) and aggregate plots of global average signal (bottom) surrounding the TSSs of all genes for reads under 100 bp (left), and for mononucleosome-sized fragments (right). For mononucleosome-sized fragments (180-250 bp), the center 3-bp nucleotide corresponding to the central dyad of the nucleosome is mapped, resulting in narrower peaks **B**. Western blot analysis of whole cell extracts of clones derived from endogenous tagging of TBP to insert the HaloTag. **C**. Time-lapse live-cell imaging of C41 HaloTBP cells stably expressing H2B-GFP as cells undergo mitosis. **D**. Live imaging versus fixed immunofluorescence for cells over-expressing Halo-TBP or the endogenous C41 Halo-TBP knock-in. **E**. Chromosome enrichment levels for either the over-expressing Halo-TBP cells under live or fixed conditions, or the C41 Halo-TBP knock-in cells under live or fixed conditions. n = 40 cells. Data are represented as mean ± SEM.

Given the persistence of chromatin accessibility at promoters of mitotic mouse ES cells, we hypothesized that the transcriptional machinery assembled at the TSS may remain associated during mitosis. TBP, a core member of the transcriptional machinery, has previously been shown to associate with certain loci during mitosis (22). Thus, we hypothesized that TBP may act as a general bookmarking factor for active genes during mitosis in mouse ES cells. We therefore set out to visualize TBP dynamics throughout the cell cycle. In order to circumvent artifacts caused by cell fixation and over-expression, we used CRISPR/Cas9 to knock-in HaloTag at the endogenous TBP locus and allow for live-cell imaging. We generated several cell lines containing heterozygous and homozygous tagged Halo-TBP, as determined by Western blot (Figure 1C). TBP is an essential gene, and so the ability of the homozygous Halo-TBP ES cell line to generate all three germ layers in a teratoma assay assured us that pluripotency is maintained, and that tagged TBP is functioning normally (Figure 1 - Figure Supplement 3). After stable integration of H2B-GFP in the Halo-TBP C41 homozygous line, we performed two-color time-lapse imaging of H2B-GFP and Halo-TBP as mouse ES cells undergo mitosis. We found that the endogenous Halo-TBP is enriched on mitotic chromosomes throughout mitosis (Figure 1D).

Several studies have previously reported that concentrations of TBP within the cell are highly regulated (21, 25), which questions previous experiments examining the dynamics of TBP association with mitotic chromosomes using over-expressed fluorescently-tagged proteins. We thus tested whether over-expressed Halo-TBP could recapitulate the behavior of endogenous Halo-TBP by comparing the C41 knock-in line with a stable cell line containing exogenous over-expressed Halo-TBP. We analyzed the enrichment on mitotic chromosomes of the over-expressed (Halo-TBP OE) and the endogenous knock-in (Halo-TBP KI) in live versus fixed cells as previously described (11). We found that Halo-TBP OE cells showed the artifactual exclusion from mitotic chromosomes following formaldehyde fixation whereas the knock-in cells maintained enrichment even after fixation (Figure 1E,F). These results suggest that, unlike most sequence specific TFs that are susceptible to fixation-induced artifacts, the endogenous TBP does not become evicted from mitotic chromosomes after fixation, suggesting a distinct interaction modality. In contrast, over-expressed exogenous TBP is susceptible to fixation-induced artifacts and therefore does not faithfully recapitulate the endogenous behavior.

### TBP stably binds to mitotic chromosomes

We next asked how TBP interaction with mitotic chromosomes compares to its interaction with interphase chromatin. We imaged individual Halo-TBP molecules at a high camera speed (133 Hz) and high illumination intensity using stroboscopic photoactivatable single particle tracking (spaSPT) in order to simultaneously track both fast-diffusing and chromatin-bound molecules (26, 27). We then localized individual particles and measured their displacement between consecutive frames (Figure 2 - Figure Supplement 1). Figure 2A shows the distribution of displacements after 3 frames (Δ*τ* = 22.5 ms) for cells in interphase and in mitosis. The shift to longer displacements in mitosis compared to interphase suggests a difference in the bound fraction and/or the diffusion constant between the two cell cycle stages. This difference becomes more apparent when the cumulative distribution function (CDF) is plotted for displacements after 3 frames (Δ*τ* = 22.5 ms) for both interphase and mitotic cells (Figure 2B), which shows a rightward shift in the CDF for mitotic cells. We then used SPOT-ON, an analysis tool for kinetic modeling of SPT data, and fitted the distribution of displacements with a 2-state kinetic model representing ‘bound’ and ‘free’ populations (Figure 2C) (27). Such a model gives estimates of the fraction of molecules in each state, and the corresponding apparent diffusion constants for each population.

**Figure 2.**
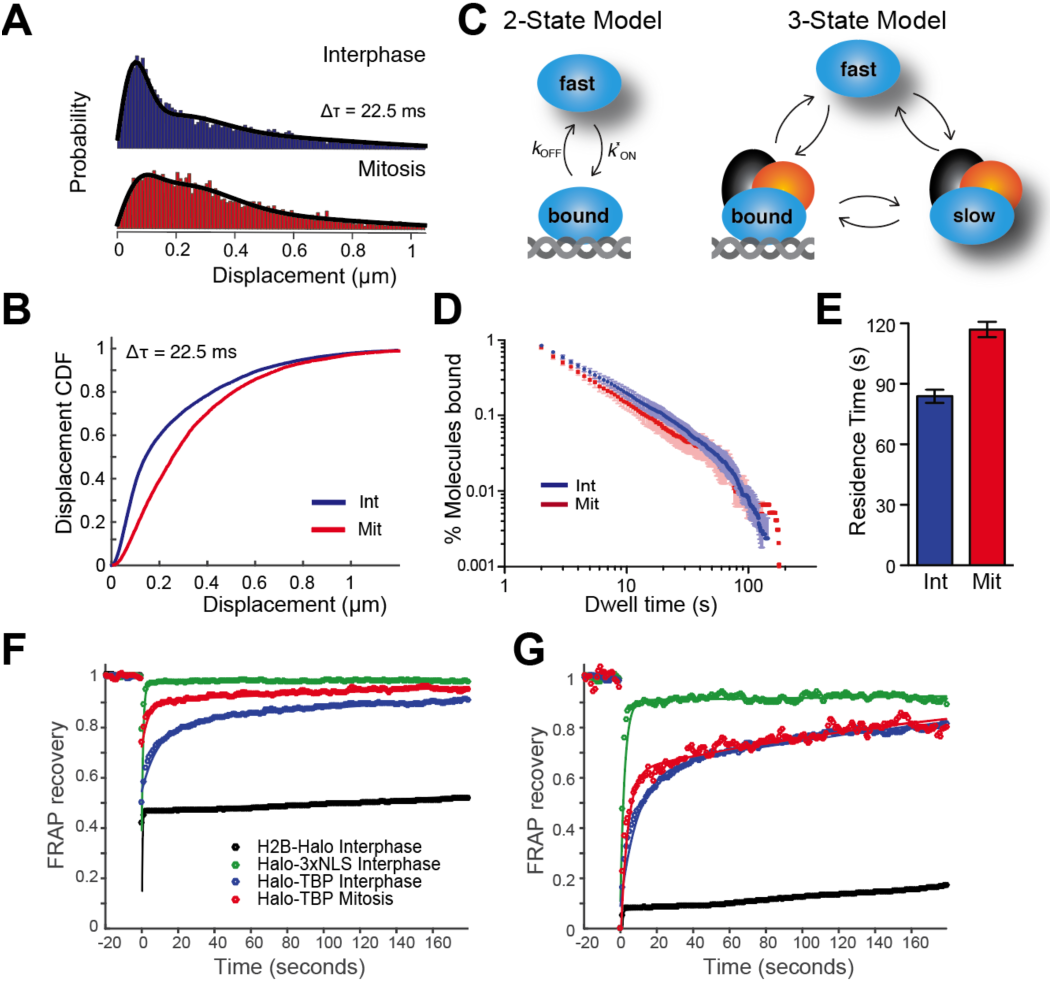
TBP dynamics within live cells. **A**. Jump length histogram measured in displacements (μm) after 3 consecutive frames (Δ*τ* = 22.5 ms) during spaSPT for cells in interphase (blue) or in mitosis (red). **B**. Cumulative distribution function of displacements after 3 frames (Δ*τ* = 22.5 ms) during spaSPT for cells in interphase (blue) or in mitosis (red). **C**. Depiction of different states for the 2-state or the 3-state kinetic model. For the 2-state model, the molecules switch from fast diffusing mode to bound states. In the 3-state model, the freely diffusing molecules can be divided into two categories, fast and slow, which can switch to a DNA-bound state. **D**. Dwell time histogram of the fraction of endogenously-tagged Halo-TBP molecules remaining bound for interphase (blue) and mitotic (red) cells. **E**. Quantification of the residence time of Halo-TBP in interphase (blue) and mitotic (red) cells. n = 30 cells. **F**. Quantification of fluorescence recovery at the bleach spot for the indicated Halo-tagged construct in interphase and mitosis. n = 30 cells. **G**. Data from F, normalized for bleach depth. Data are represented as mean ± SEM.

The resulting fit for the 2-state model shows that the fraction of bound TBP decreases by half, from 34.6% in interphase to 18.6% in mitosis (Table 1). However, upon closer inspection, we find that the 2-state model fitted the data poorly, showing large residuals after the fit (Figure 2 - Figure Supplement 2). This result suggests that a dual classification of ‘bound’ and ‘free’ TBP populations may be insufficient to accurately characterize the data. Therefore, we fitted a 3-state kinetic model to represent three distinct populations: a fast-diffusing TBP population (’fast‘), a slower diffusing population wherein TBP may be diffusing in a complex with the TAFs as TFIID (’slow‘), and a stably DNA bound population (’bound‘) (Figure 2C). The 3-state model fitted the data more precisely (Figure 2A, black line), showing reduced residuals after fitting (Figure 2 - Figure Supplement 2 and 3). From this analysis, we estimate that in interphase 38% of TBP molecules are in fast diffusion mode, whereas 37.2% are found in a slower 3D diffusing complex (Table 1). Relatedly, we find that 24.8% of TBP molecules are bound to interphase chromatin (Table 1). During mitosis, the chromatin bound fraction decreases to 12.3%, with concomitant increases in both the slow and fast diffusing populations (46.5% and 41.2%, respectively) (Table 1). This decrease in fraction bound during mitosis may reflect the minimum amount of TBP needed to ‘bookmark’ active genes during mitosis. Furthermore, measured diffusion constants for TBP molecules that are in fast (Apparent *D*_fast_) or in slow diffusion (Apparent *D*_slow_) show no significant difference between interphase and mitotic cells (Table 1), suggesting that although the fraction of TBP in each subpopulation is different in mitosis, the diffusion of each subpopulation remains the same.

Given the decreased percent of TBP molecules that are bound during mitosis, we next asked how stable is TBP binding and if the mitotic binding dynamics differ from those of the bound population in interphase. We previously discovered that prototypic sequence-specific TFs such as Sox2 display a more dynamic binding behavior during mitosis compared to interphase, showing about 50% decrease in residence time during mitosis that is largely due to the global decrease in transcriptional activity (11). To determine if bound TBP molecules behave similarly to sequence-specific TFs, we performed single particle tracking (SPT) with long exposure times (500 ms) as previously described (28). This imaging technique allows for a ‘blurring out’ of diffusing molecules while bound ones appear as diffraction-limited spots. After localization and tracking of individual bound molecules, we measured the dwell time, the amount of time each molecule remains detected, and plotted the log-log histogram of dwell times for interphase and mitotic cells (Figure 2D). In contrast to sequence-specific TFs like Sox2, TBP remains as stably bound on mitotic chromosomes as on interphase chromatin. We then fitted a two-component exponential decay model, representing specific versus non-specific binding events, to the dwell time histograms and extracted residence times for the two bound populations (29). After correcting for photobleaching, the residence time for specific Halo-TBP binding during interphase is on the order of minutes (Figure 2E). This binding is significantly more stable than typical sequence-specific TFs like Sox2, p53, GR, which normally have residence times on the order of a few to around ten seconds in interphase (29–33). Surprisingly, some Halo-TBP also remains stably bound during mitosis (Figure 2E), with a calculated average residence time of 118 seconds. This apparent TBP residence time in mitosis is even longer than in interphase. However, based on the dwell time histogram, this is likely due to the contribution of a small fraction of TBP molecules (less than 1%) that appear very stably bound (dwell time above 100 s in Figure 2D). Given that the whole distribution profiles appear very similar, we concluded that for most of the bound TBP, residence times are indistinguishable between interphase and mitotic cells. This result is in stark contrast to Sox2, whose residence time decreases by at least half during mitosis, suggesting that TBP binding to mitotic chromosomes may be independent of transcriptional activity.

As an orthogonal approach to SPT, we performed FRAP (Fluorescence Recovery after Photobleaching) analysis to independently measure TBP binding dynamics during interphase and mitosis. We plotted the FRAP recovery at the bleach spot over time for Halo-TBP in interphase and in mitotic cells, along with Halo-3xNLS and H2B-Halo (Figure 2F). When normalized for bleach depth, the recovery curves for interphase and mitotic cells become super-imposable (Figure 2G). The difference in bleach depth may indicate a difference in the fraction of bound molecules between interphase and mitotic cells, which would be consistent with our SPT results (Table 1; 24.8% vs. 12.3% bound in interphase and mitosis, respectively), whereas the super-imposable recovery curves would be consistent with similar residence times of the bound fraction. This analysis underscores the need for multiple orthogonal measurements to better interpret TF dynamics.

### TBP binds to the TSS of active genes in mitosis

We next asked whether TBP binds to specific sites in the genome during mitosis. Given the relatively stable binding of TBP during mitosis and its resistance to the formaldehyde-based exclusion artifact, we reasoned that endogenous TBP could be amenable to formaldehyde-based ChIP-seq analysis without complications from incurring cross-linking artifacts. Indeed we were able to perform ChIP-seq on both asynchronous and mitotic cells in two biological replicates that are highly reproducible (Figure 3 - Figure Supplement 1). Genome browser visualization of mapped reads for each replicate shows that binding profiles are highly reproducible, and that TBP is maintained at the same genomic sites in mitosis as in interphase chromatin (Figure 3A). We next analyzed the level of TBP enrichment at the TSS of all genes and displayed the ChIP-seq signal on a 4 kb region centered at the TSS as a heatmap, with genes arranged by decreasing expression in the asynchronous population (Figure 3B). The heat map shows that TBP enrichment at the TSS is correlated with expression level both in the asynchronous and mitotic population. Averaging the enrichment across all TSSs shows that enrichment in mitosis is roughly 80% of the enrichment in asynchronous cells (Figure 3C). This reduced enrichment at the TSS during mitosis likely reflects the reduced fraction of molecules that are bound during mitosis rather than the strength of binding (Figure 2C). To determine whether TBP binds more to certain classes of genes in mitosis, we performed unbiased k-means clustering with k = 6 and displayed the clusters as heatmaps centered at the TSS (Figure 3D). This clustering analysis shows 3 main groups. Group 1 includes the first 2 clusters, which show increased TBP binding in mitosis relative to asynchronous cells. Group 2 consists of genes that display no change or very little TBP signal at the TSS. Lastly, group 3 is composed of the last three clusters that show decreased TBP signal in mitosis. This analysis suggests that TBP binds differentially to specific genes depending on cell cycle phase. However, when we analyzed the types of genes within each group using gene ontology (GO) term analysis, we find that groups 1 and 3 are enriched for similar classes of genes, primarily ones involved in the cell cycle (Figure 3E). This analysis suggests that the differences in TBP binding that we observe by ChIP-seq largely reflect the cyclical expression of genes during the cell cycle. Despite the variations in TBP levels, most active genes retain TBP binding in mitosis, suggesting a global bookmarking activity by TBP.

**Figure 3.**
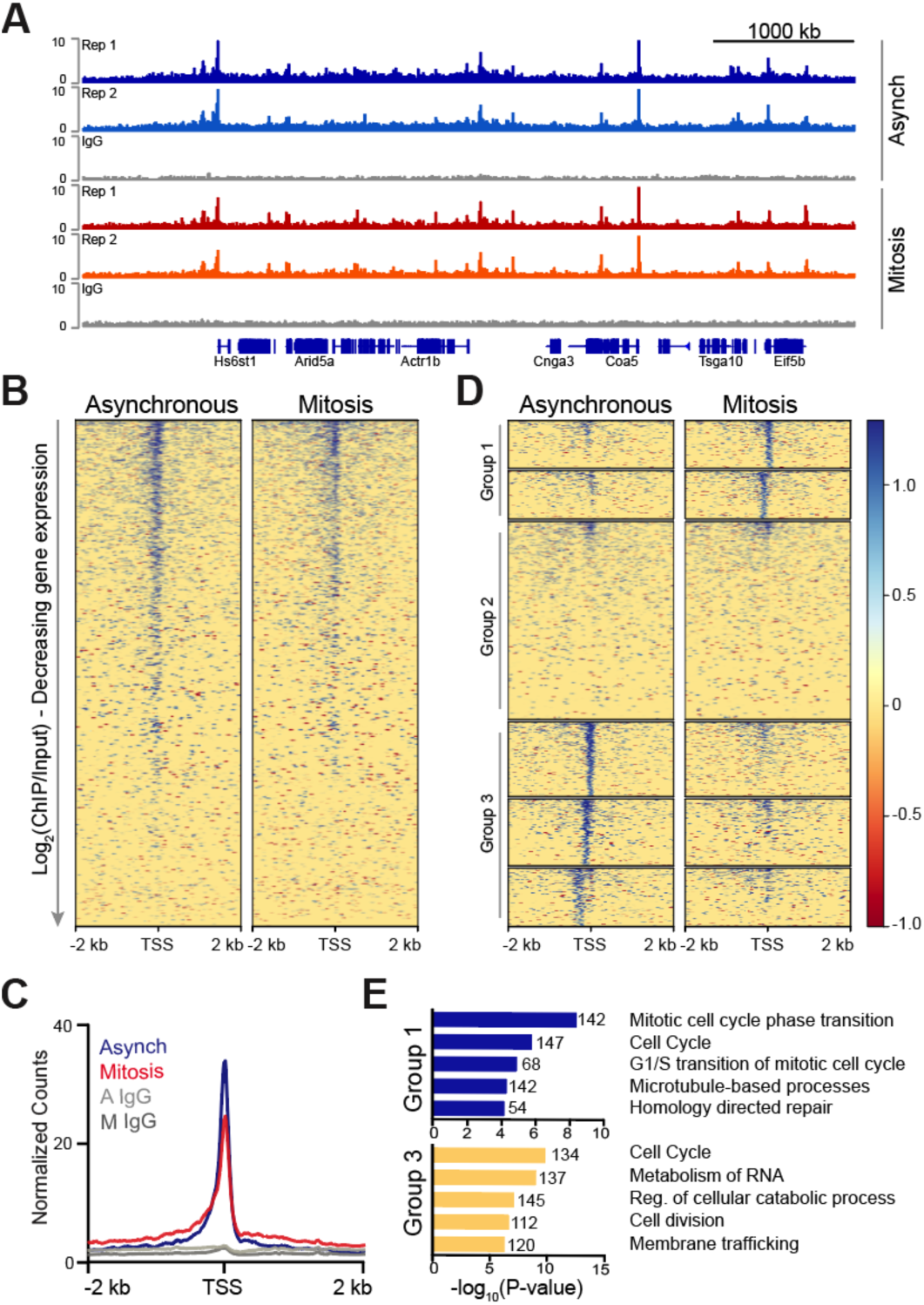
TBP maintains binding to promoters of active genes during mitosis. **A**. Genome browser profiles for TBP ChIP-seq replicates in asynchronous and mitotic cells and the corresponding IgG ChIP control. **B.** Heatmaps of the log_2_ ratio of ChIP over the input signal surrounding the TSS of all genes, arranged by decreasing gene expression for asynchronous and mitotic samples. **C.** Average ChIP-seq read density for all TSS and surrounding regions for asynchronous (blue) and mitotic (red) samples and corresponding IgG controls (grays). **D.** Unbiased k-means clustering of data from B with k = 6. Clusters are grouped into 3 groups depending on changes in signal. **E.** Gene Ontology term analysis of genes in Group 1 and Group 3 from **D**. Numbers correspond to the number of genes within the group that is labeled with the specific GO term.

### TBP recruits RNA Polymerase II to mitotic chromosomes

Having determined that TBP stably binds to the TSS of active genes during mitosis, we next asked what is the function of such stable TBP mitotic binding. To address this question, we endogenously knocked-in the plant-specific minimal auxin-inducible degron (mAID) (34, 35) at the TBP locus to generate homozygous mAID-TBP fusion proteins (C94 cell line). This fusion enables acute, auxin-inducible degradation of the mAID-tagged TBP via the ubiquitin degradation pathway (Figure 4A). In fact, within 6 hours of IAA (auxin) treatment, only about 3% of mAID-TBP molecules remains detectable by bulk protein analysis with Western blot (Figure 4B), and by immunofluorescence using *α*-TBP antibody (Figure 4C). This drug-inducible degradation of TBP provides us with a robust tool to causally test the functions of TBP binding specifically during mitosis.

**Figure 4.**
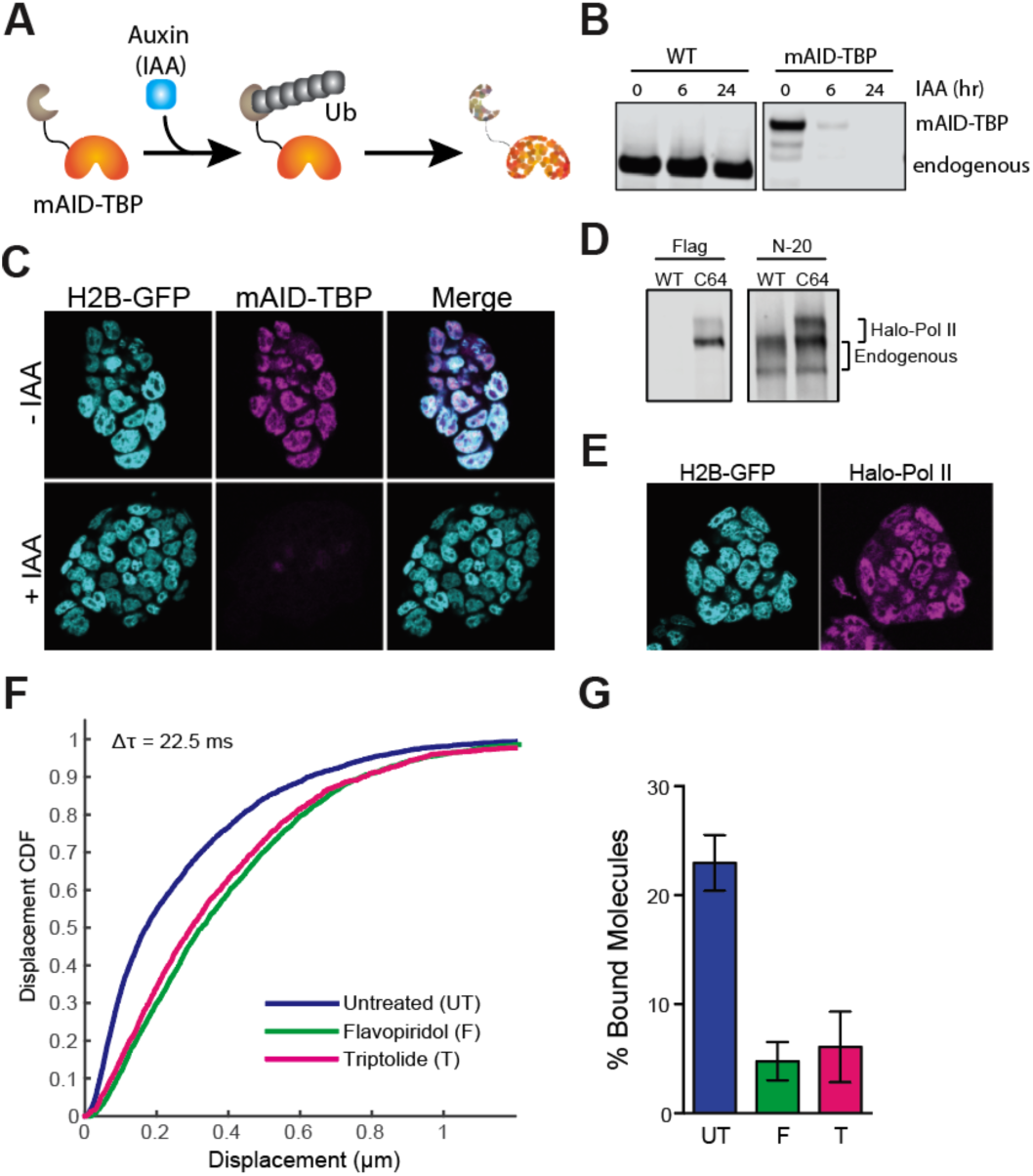
Drug-inducible degradation of endogenous TBP. **A**. Schematic for mechanism of auxin-inducible degradation. Ub, ubiquitin. **B**. Western blot analysis for time-course of Auxin (IAA)-dependent degradation of WT cells or cells with the endogenous knock-in of the mAID to the TBP locus. **C**. Immunofluorescence using *α*-TBP of cells with endogenous mAID-TBP and stably expressing H2B-GFP without IAA (top) or after 6 hours of IAA treatment (bottom). **D**. Western blot analysis of wild-type (WT) ES cells or Halo-Pol II knock-in (C64) using *α*-Flag to detect the Halo-Flag knock-in and *α*-N-20, an antibody against the N-terminal of Pol II, to detect all Pol II levels. The unphosphorylated and phosphorylated bands for endogenous and Halo-Pol II are marked. **E**. Live imaging of Halo-Pol II showing nuclear localization as marked with H2B-GFP. **F.** Cumulative distribution function (CDF) of Halo-Pol II displacements for three consecutive frames (Δ*τ* = 22.5 ms) for untreated cells, or cells treated with 30-60 min of Flavopiridol or Triptolide. **G.** A 3-state model used to fit the CDF in **F** and the fraction of bound Halo-Pol II was extracted and plotted for each condition. UT, untreated. F, Flavopiridol. T, Triptolide. n = 30 cells and data are represented as mean ± SEM.

A primary function of TBP at the TSS of active genes is to recruit Pol II and form the pre-initiation complex (PIC) (12, 36). Our discovery here of stable TBP binding at TSSs during mitosis led us to hypothesize that TBP may function similarly during mitosis and recruit Pol II to active genes, despite the extensive decrease in global transcriptional activity. To test this hypothesis, we endogenously knocked-in a HaloTag at the Rpb1 locus, the catalytic subunit of RNA Polymerase II, in the mAID-TBP (C94) cells. We successfully produced heterozygous knock-in of Halo-Pol2 II, as shown by Western blot analysis (Figure 4D), and the tagged protein localizes within the nucleus as expected (Figure 4E). We further assessed the functionality of the tagged Pol II with several experiments. First, we used spaSPT to analyze the distribution of Pol II displacements before and after transcription inhibition by Flavopiridol (elongation inhibitor) and Triptolide (initiation inhibitor) (Figure 4F). We found that Halo-Pol II showed a significant shift in displacements towards longer jumps, indicative of increased diffusing molecules and decreased bound ones. When fitting the data with a 2-state model, we found that the fraction of bound Pol II decreased by 80% after drug treatments (Figure 4G). These results suggest that the halo-tagged Pol II is sensitive to drug inhibition. Furthermore, ChIP-qPCR analysis using antibodies against Pol II (targeting total Pol II) and Flag (located between Halo and the N-terminal of Rpb1, targeting only Halo-Pol II) showed that the tagged Pol II binds to regions where the endogenous Pol II binds (Figure 4 - Figure Supplement 1), altogether confirming that the tagged Pol II remains functional.

To test the role of TBP in recruiting Pol II, we next performed spaSPT on Halo-Pol II in interphase and mitosis, with and without TBP. The distribution of Halo-Pol II displacements shows an increase in longer displacements between consecutive frames after TBP degradation both in interphase and mitotic cells (Figure 5 - Figure Supplement 1). This effect is evident in the rightward shift of the displacement CDF after TBP degradation for both interphase and mitotic cells (Figure 5A). We then fitted a 2-state kinetic model as before to represent ‘bound’ and ‘free’ populations (Figure 2C) and extracted estimates of the fraction of molecules and corresponding diffusion constants for each population (Table 2). However, as we had found with TBP, the 2-state kinetic model fitted the Pol II data poorly, particularly for mitotic cells, as evident by the large residuals after fitting (Figure 5 - Figure Supplement 2-5), suggesting that this model is insufficient to characterize Pol II dynamics. We then fitted a 3-state kinetic model (Supplementary Figure 4) and extracted the fraction of Halo-Pol II molecules in each state for all the tested conditions (Table 2). We note that despite differences in absolute numbers, the global trend remains similar in extracted parameters between the 2-state and 3-state models, and so conclusions remain the same.

**Figure 5.**
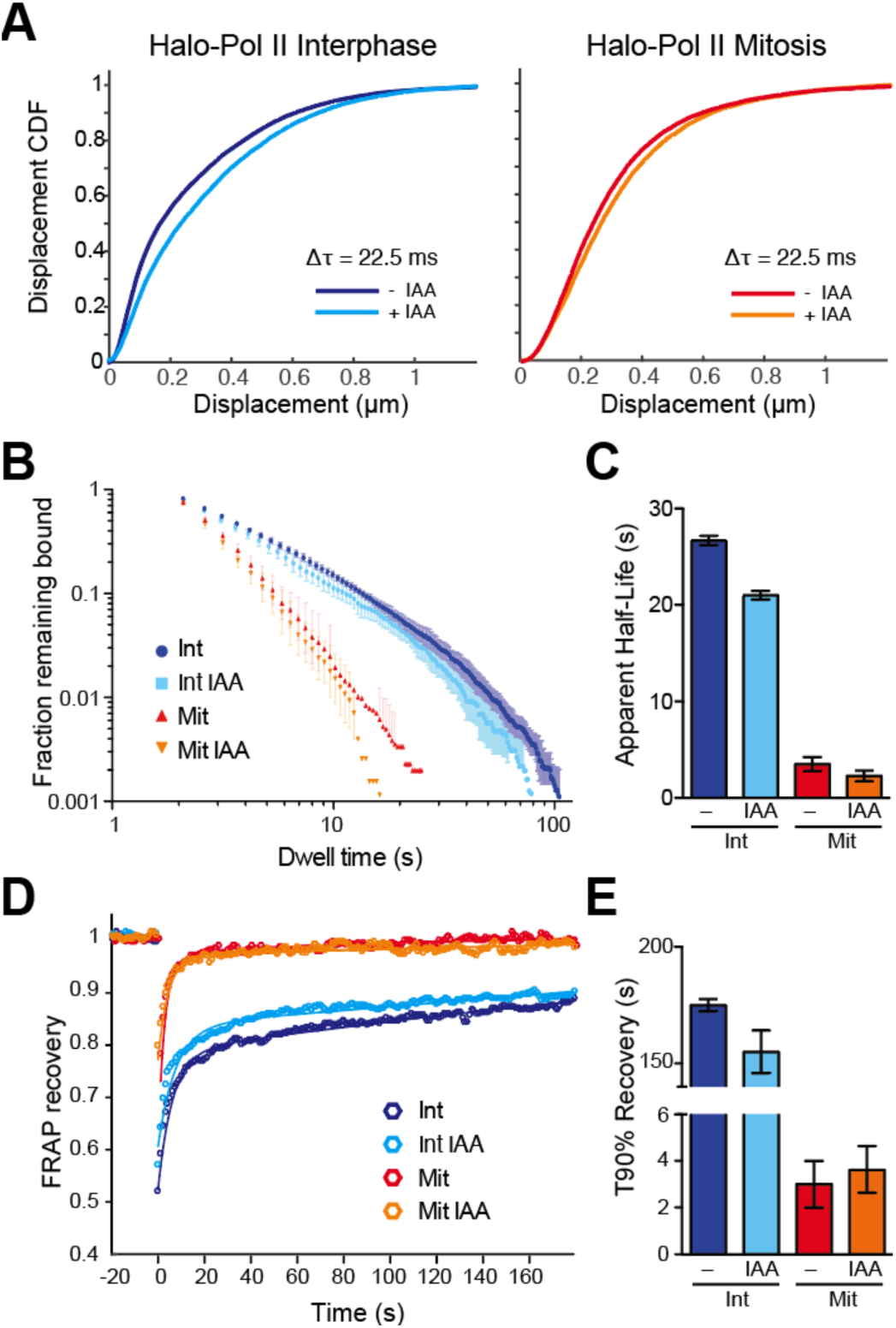
TBP-dependent dynamics of Pol II in live cells. **A**. Cumulative distribution function of displacements after 3 frames (Δ*τ* = 22.5 ms) during spaSPT of endogenous Halo-Pol II for cells in interphase (left) or in mitosis (right) either with or without IAA treatment (TBP degradation). **B.** Dwell time histogram of the fraction of endogenously-tagged Halo-Pol II molecules remaining bound for interphase (blues) and mitotic (reds) cells either with or without IAA treatment, as indicated in legend. **C**. Quantification of the apparent half-life (see Methods) in seconds of Halo-Pol II in interphase (blues) and mitotic (reds) cells. n = 30 cells. **D**. Quantification of fluorescence recovery at the bleach spot for Halo-Pol II in interphase (blues) and mitosis (reds), either with or without IAA treatment. n = 30 cells. **E**. from **D**, the average time to reach 90% recovery for Halo-Pol II in interphase (blues) or mitosis (reds), either with or without IAA treatment. Data are represented as mean ± SEM.

During interphase, 32% of Pol II is fast diffusing whereas 45.6% is slow diffusing. The nature or composition of the two distinctly diffusing population remains unknown, but what is clear is that the fraction of Pol II that is bound on chromatin is 22.4%. This bound population decreases by half to 12.1% during mitosis, revealing a small but significant population of Pol II that persists on mitotic chromosomes. We next focused on the effect of TBP degradation on Pol II recruitment and binding. Upon TBP degradation in interphase cells, the fraction of bound Halo-Pol II decreases from 22.4% to 16.1% (Table 2), consistent with the important role of TBP in recruiting Pol II to active genes. TBP degradation during mitosis also led to a decrease in the fraction of bound Pol II, from 12.1% in untreated mitotic cells to about 10.8% after TBP degradation. These results suggest that TBP recruitment of Pol II to DNA is independent of the cell cycle or the chromatin condensation state.

We next examined the dynamics of Pol II binding to mitotic chromosomes by performing SPT with long exposure times (500 ms). We measured the dwell time of diffraction-limited bound molecules and plotted the log-log histogram of dwell times for Halo-Pol II in interphase and mitosis with and without TBP (Figure 5B). The most obvious difference is seen between interphase and mitotic cells, with much reduced dwell times of Halo-Pol II during mitosis compared to interphase cells. This decrease is consistent with the global decrease in active transcription during mitosis. After degrading TBP, Halo-Pol II shows decreased dwell time in interphase, consistent with the role of TBP in initiating transcription. In contrast, TBP degradation shows less effect on the dwell time of Pol II during mitosis. We then fitted a two-state exponential decay model to the dwell times to extract the apparent *k*_*off*_ of Pol II specific binding. Pol II binding to DNA involves complex steps, from transient binding due to promoter association and abortive initiation, to the stable binding during elongation, and to the as yet undefined mechanisms for termination. Therefore, modeling the binding of Pol II to DNA using a single residence time likely over-simplifies the complex dynamics of Pol II progression during transcription. With this caution in mind, instead of extracting a residence time for Pol II, we calculated the apparent half-life of the bound population. During interphase, the apparent half-life of Pol II is 26.7 seconds (Figure 5C). TBP degradation leads to a modest decrease in apparent half-life, to 21.0 seconds. This modest decrease is consistent with the prominent role of TBP in promoter association, which is largely composed of transient Pol II binding. During mitosis, the apparent half-life of Pol II binding decreases to 3.5 seconds. This order of magnitude decrease in apparent half-life suggests that the mechanisms that stabilize Pol II binding becomes dramatically decreased during mitosis. Furthermore, after TBP degradation during mitosis, the apparent half-life decreases from 3.5 to 2.3 seconds, consistent with the role of TBP in recruiting transiently-binding Pol II to mitotic chromosomes.

To cross-validate the SPT residence time analysis, we performed FRAP analysis on Halo-Pol II during interphase and mitosis, with and without TBP degradation. We plotted the FRAP recovery over time (Figure 5D) and measured the time it takes to reach 90% recovery (T90%) (Figure 5E). As with the SPT, the most drastic difference in the FRAP curves is seen between interphase and mitotic cells under normal TBP levels. Halo-Pol II takes 179 seconds to reach 90% recovery in interphase cells, whereas in mitosis, it takes 3 seconds. Upon TBP degradation in interphase cells, T90% decreases to 150 seconds, consistent with the decreased apparent half-life as measured by SPT. TBP degradation in mitotic cells shows little change in the recovery curve relative to untreated mitotic cells, further confirming our SPT residence time analysis. Taken together with the SPT results, these findings suggest that TBP affects the more transient promoter association of Pol II during interphase, and potentially directs Pol II binding to mitotic chromosomes.

### Mitotic TBP binding affects the rate of transcriptional reactivation

We next asked how TBP binding during mitosis affects transcription reactivation following cell division. Having shown that TBP promotes Pol II recruitment to mitotic chromosomes, we hypothesized that this recruitment helps prime active genes for efficient reactivation following mitosis. To test this hypothesis, we performed two orthogonal experiments, one based on imaging of single cells, and another based on sequencing newly transcribed RNA products as a time course after mitosis. First, we returned to our Halo-Pol II C64 cells and performed spaSPT. We marked cells at the metaphase stage of mitosis and imaged the same cells every 15 minutes until 60 minutes after first staging, following cells as they exited mitosis and entered G1. To ensure that repeated imaging had no significant effect on Pol II behavior, we performed the same imaging regimen on various interphase cells and calculated the fraction of molecules bound over time. This population of Pol II remained stable over the course of the imaging regimen even for cells treated with auxin for TBP degradation, suggesting that repeated imaging did not adversely affect Pol II behavior (Figure 6A). As expected, during the progression from mitosis into G1, the fraction of bound Pol II molecules steadily increased over time, consistent with increasing activation of transcription following mitosis (Figure 6B). When we performed the same experiment with cells depleted of TBP, we observed that the fraction of bound Pol II remained flat over time (Figure 6B), suggesting that TBP is required for re-establishing the transcription program following mitosis.

**Figure 6.**
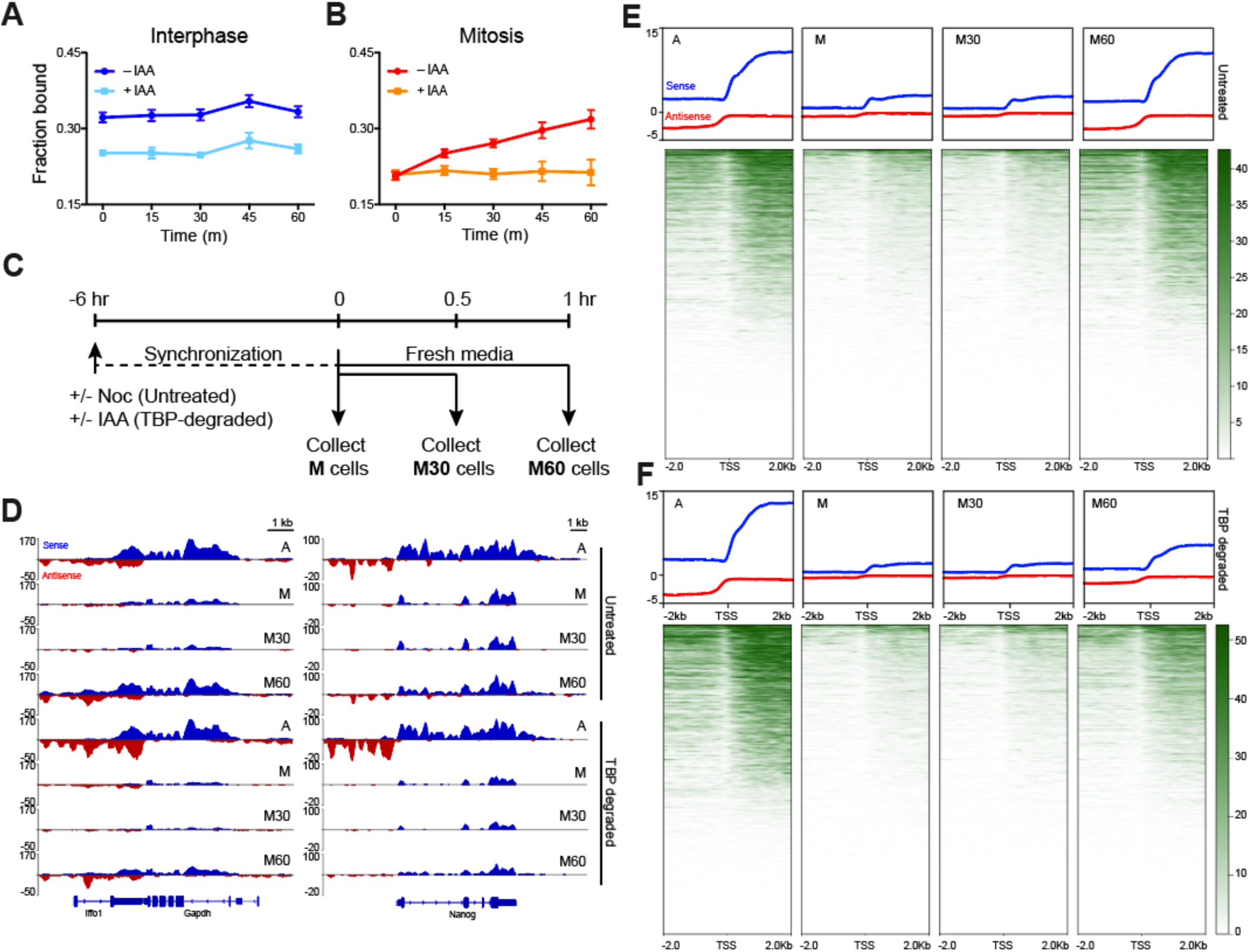
TBP recruits Pol II during mitosis. **A**. The fraction of bound Halo-Pol II molecules as a function of time (t = 0, release from DMSO/IAA treatment) under no IAA (WT conditions) or after 6 hours of IAA treatment (teal) for interphase cells. **B**. Same as **A**, but for mitotic cells. **C**. Schematic of time course regimen for extracting chromatin-associated nascent RNA. Asynchronous cells are treated with DMSO or IAA for 6 hours prior to chromatin-associated nascent RNA extraction (not depicted). For other samples, cells are synchronized with Nocodazole for 6 hours, and treated with DMSO (untreated) or IAA (TBP-degraded) during Nocodazole synchronization. For mitotic samples (M), cells are immediately collected. For time course after mitosis, synchronized M cells (untreated or TBP-degraded) are replaced into fresh media to release from mitotic arrest and TBP degradation and are collected either after 30 minutes (M30) or 60 minutes (M60) following fresh media resuspension. **D**. Extracted chromatin-associated nascent RNA were sequenced in strand-specific manner, and the sense (blue) and anti-sense (red) reads are plotted for Gapdh (left) and Nanog (right) loci for all corresponding samples. **E**. Genome-wide average plots for all TSS and surrounding regions for sense (blue) and anti-sense (red) reads for each indicated untreated sample, and the corresponding heatmaps for all reads (bottom). **F**. Same as E but for TBP-degraded samples.

We also performed a global analysis of newly transcribed RNA as a function of time following mitosis. To assess changes in newly transcribed RNA, we performed successive biochemical fractionation and extracted the nascent chromatin-associated RNA (chr-RNA) of asynchronous cells (A), mitotic cells synchronized with Nocodazole (M), and 30 min and 60 min after Nocodazole release (M30 and M60, respectively) of synchronized mitotic cells in two replicates (Figure 6C). We then performed strand-specific sequencing of the extracted chr-RNA along with ERCC spike-in controls and mapped the reads to the mouse genome (Figure 6 - Figure Supplement 1 and 2). A snapshot of the mapped reads on a housekeeping gene (*Gapdh*) and an ES cell-specific gene (*Nanog*) is shown on Figure 6D. The high proportion of sequenced reads in intronic regions in asynchronous samples (Figure 6D, Figure 6 - Figure Supplement 1) confirms successful enrichment for newly synthesized RNAs. To assess transcriptional activity through mitosis on a global scale, we averaged the normalized signal in a 4 kb region surrounding the TSS for all genes in each sample (Figure 6E). Overall levels decrease in Nocodazole-arrested M samples and remain low 30 minutes after release. However, by 60 minutes after release, RNA levels have increased to the levels in asynchronous samples. These data are consistent with the global decrease in transcriptional activity followed by reactivation as cells exit mitosis.

To examine the role of TBP in promoting efficient reactivation, we degraded TBP in asynchronous samples (A) and in Nocodazole arrested mitotic cells (M), and collected cells after 30 or 60 minutes after mitosis as before (Figure 6C, Figure 6 - Figure Supplement 1). As with untreated samples, we extracted and sequenced the chr-RNA and ERCC spike-in controls in a strand-specific manner (Figure 6 - Figure Supplement 1 and 2). A snapshot of the mapped reads of TBP-degraded samples on *Gapdh* and *Nanog* is also shown on Figure 6D. We still observe high levels of intronic reads in asynchronous (A) samples, despite near complete degradation of TBP. Our results are reminiscent of a previous study that has observed Pol II transcription in the absence of TBP in mouse blastocyst cells (14). However, we observe a marked decrease in transcription reactivation in M60 samples following TBP degradation in mitosis (Figure 6D). This decrease occurs globally as most genes show decreased nascent chr-RNA levels (Figure 6F), suggesting a specific role for TBP in transcriptional reactivation following mitosis.

We next examined the kinetics of transcriptional reactivation following mitosis by taking the log_2_ ratio of reads in the 30-and 60-minute conditions relative to mitotically arrested (M30/M and M60/M, respectively), for both untreated and TBP-degraded samples. In this way, we can observe the change in RNA levels relative to mitotic cells as a function of time after release from mitosis. Globally, the untreated M30/M sample shows no overall change in RNA levels whereas M60/M samples show massive increase in both upstream and downstream transcription at the TSS (Figure 7 - Figure Supplement 1). On the other end, TBP-degraded samples show delayed transcription levels evident in the M60/M sample (Figure 7 - Figure Supplement 1). To determine whether certain groups of genes behave differently from the global mean, we performed unbiased k-means clustering on the untreated M30/M data with k = 3. Cluster 1 includes 5504 genes that show a mild increase in transcription whereas cluster 3 includes 6693 genes that show a mild decrease in transcription relative to mitotic cells. The remaining genes (cluster 2) show no change in transcription (Figure 7A,B). Ordering the M60/M data using the same three clusters shows that cluster 1 increases in transcription the earliest and the fastest whereas clusters 2 and 3 lag behind. We then ordered the M30/M and M60/M data from TBP-degraded samples (Figure A,B). This analysis shows that the early changes in transcription seen in untreated samples are dampened in TBP-degraded samples. Furthermore, TBP degradation has the biggest effect on cluster 1. Using gene ontology (GO) term analysis, we find that cluster 1 is enriched for genes involved in metabolism of RNA, cell cycle, and transcription factor activity (Figure 7C). These genes are also generally highly expressed in the asynchronous population (Figure 7 - Figure Supplement 1), suggesting that these genes represent the global transcription program of the cells. In contrast, cluster 3 is enriched for genes involved in basic cellular processes such as microtubule cytoskeleton organization and regulation of GTPase activity, but also for genes involved in differentiation and development such as cell morphogenesis in differentiation, and morphogenesis of epithelium (Figure 7C). Furthermore, these genes tend to be less expressed than cluster 1 in the asynchronous population (Figure 7 - Figure Supplement 1). These data strongly suggest that TBP promotes efficient reactivation of the global ES cell transcription program and thus plays a functional role in promoting the maintenance of cell state through the cell cycle.

**Figure 7.**
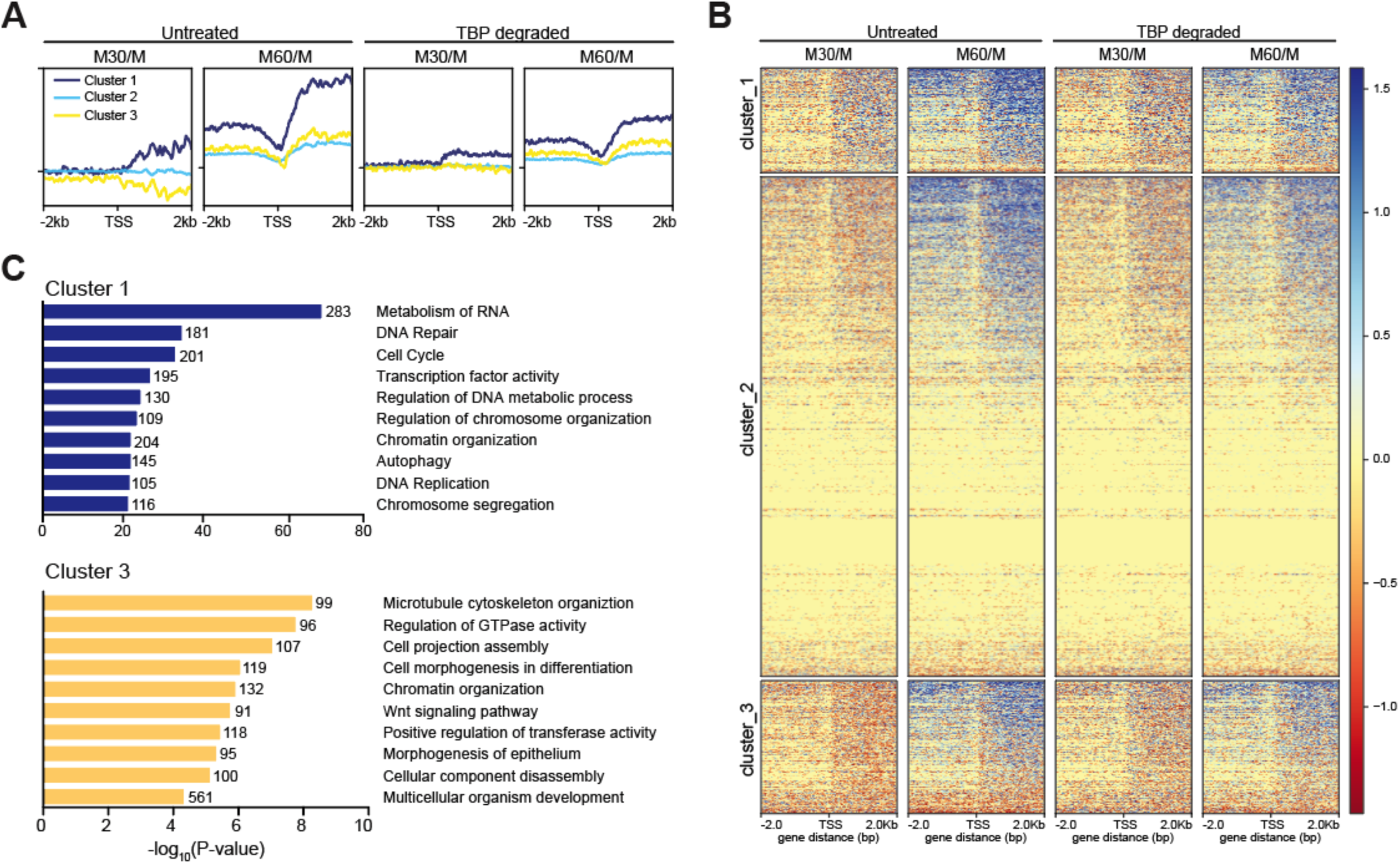
TBP promotes efficient reaction of transcription following mitosis. **A.** The log_2_ ratio of M30 and M60 reads relative to M reads were calculated for untreated and TBP-degraded samples (M30/M and M60/M, respectively). The M30/M samples were clustered using k-means clustering with k = 3, and the rest of the samples are ordered by this clustering, and average plots surrounding the TSS are shown for each cluster. **B.** Heat map analysis of each cluster for each sample in **A. C.** GO term analysis for genes present in cluster 1 (top) and cluster 3 (bottom). Numbers correspond to the number of genes within the custer that is labeled with the specific GO term.

## Discussion

Although different mechanisms exist to ensure faithful transmission of transcription programs from mother to daughter cells afer mitosis, the precise role of transcription factors in this process has remained elusive. In this study, we have shown that unlike most classical transcription factors that bind with rapid dynamics, the endogenous TBP maintains stable binding at the TSS of active genes in mitotic mouse ES cells. Remarkably, such stable TBP binding leads to recruitment of Pol II to mitotic chromosomes, despite the general inhibition of transcriptional activity. Rather, as cells exit mitosis, TBP promotes efficient reactivation of transcription globally by priming the promoters of active genes. Taken together, TBP acts as a stable bookmarker of active promoters throughout mitosis and promotes transcriptional fidelity from mother to daughter ES cells to maintain the ES cell identity.

Previous biochemical and structural data have generated static views of molecular processes, such as the formation of stable complexes or binding of factors to DNA. These static views have propagated the sense of stability. For instance, a ChIP signal for TFs has generally been viewed as a binary classification of specific loci: bound or unbound. The advent of live cell single molecule imaging has revolutionized our understanding of basic molecular processes, particularly, how dynamic these processes are. Many TFs examined by live cell single-molecule imaging have shown much shorter residence times on DNA than previously expected from biochemical data. For instance, Bicoid binds to the Drosophila genome for about 100 ms (37), p53 has an average residence time of 3-6 seconds (38), and Sox2 binds to specific loci in mouse ES cells for about 12 seconds on average (11, 29). Rather than a binary classification, we now have a much more dynamic view of repeated and rapid binding/unbinding events that ultimately affect transcriptional output. In contrast to most sequence-specific TFs, TBP seems unusual by binding to loci on the order of minutes. Although still more dynamic than previous biochemical data had suggested, this more stable mode of TBP binding may serve a specific function unique to TBP among TFs, perhaps as a more stable landing pad for the dynamic binding of various other factors that make up the transcriptional machinery. For instance, binding of both TFIIA and TFIIB are highly dependent on prior TFIID/TBP binding, and TFIIB has been shown to bind to TBP-bound promoter DNA in a highly transient manner through in vitro single molecule imaging (16). Despite the general silencing of transcription during mitosis, this relatively stable binding of TBP is maintained, also in contrast to other sequence-specific TFs. Here we have shown that TBP recruits Pol II to mitotic chromosomes and promotes efficient reactivation of transcription following mitosis, but such stable binding in mitosis may also serve other functions. One potential alternative function may be to help maintain the unique chromatin architecture of promoters in condensed mitotic chromosomes perhaps through its stable binding or through as yet unidentified pathways. In this way, TBP may promote transcriptional fidelity through mitosis in multiple ways.

A previous study has shown that blastocyst cells from mouse embryos with homozygous deletion of the TBP locus still contain Pol II in active phosphorylation states, suggesting active transcription in the absence of TBP (14). Instead, the authors found that transcription from Pol I and Pol III are dramatically reduced without TBP, suggesting a differential dependency of the various polymerases on TBP. In this study, we have used a rapid drug-inducible depletion of TBP and measured the newly transcribed RNA after depletion. We have confirmed that transcription occurs even in the absence of TBP (Figure 6D). How does transcription occur in the absence of TBP? Several mechanisms may potentially answer this question. First, TBP-like factor (TLF, also known as TBP-related factor 2 – TRF2), a closely related protein to TBP containing a similar saddle-like core domain for DNA binding, has been proposed to substitute for TBP in certain specialized cases (39, 40). However, unlike the constant expression of TBP, previous studies have shown that TLF is not expressed in early mouse embryogenesis (14, 41). A second potential mechanism for TBP-independent transcription is that a TFIID complex lacking TBP can form and activate transcription, which has been shown in vitro (42). A third potential mechanism, and not mutually exclusive with the first two, is that independent mechanisms exist for the initiation of transcription and the subsequent re-initiation. Our finding that TBP depletion strongly inhibits re-activation of genes following mitosis is consistent with this theory. That is, the first transcription event following mitosis requires TBP, providing a rationale for TBP mitotic bookmarking, whereas subsequent transcription events throughout interphase may occur even in the absence of TBP.

Our observation that TBP recruits Pol II to mitotic chromosomes may seem surprising, given that historically, transcription has been shown to be globally shut-off during mitosis (2). Recent studies, however, are beginning to shed some nuance to this historical adage. For instance, several groups have shown that transcription transiently spikes at the later stages of mitosis, followed by an adjustment back to normal interphase levels (43, 44). One potential mechanism for such a spike in transcriptional activity could derive from our observed recruitment of Pol II by TBP to mitotic chromosomes. Combined with the lack of crosstalk between enhancers and promoters during mitosis (24), it is possible that Pol II recruitment results in promiscuous transcriptional activation of most accessible promoters immediately following mitosis. Adding more nuance, several groups have recently shown that certain genes remain expressed albeit at low levels during mitosis (45, 46). It is possible that the small fraction of Pol II that we observe to be recruited to mitotic chromosomes is transcriptionally active. If so, this activity must be quite transient, as we observe no detectable Pol II molecules that bind to mitotic chromosomes for longer than 30 seconds (Figure 5C) by slow tracking SPT. In contrast, we observe much longer residence times of Pol II during interphase when transcription is highly active. Indeed, less than 1% of mitotically bound Pol II molecules show binding for greater than 10 seconds (Figure 5B), which may reflect the low transcriptional activity during mitosis as previously reported (45, 46). What does become evident, however, is the gradual increase in bound Pol II concomitant with increase in newly transcribed RNA as cells progress from mitosis, and the important role of TBP in this process (Figure 6). Regardless of the transcriptional activity of Pol II during mitosis per se, TBP promotes efficient reactivation of global transcription following mitosis, which is necessary for faithful transmission of transcription programs through the cell cycle.

## Materials and Methods

### Cell culture

For all experiments, we used the mouse ES cell line JM8.N4 (RRID: CVCL_J962) obtained from the KOMP repository (https://www.komp.org/pdf.php?cloneID=8669) and tested negative for mycoplasma. ES cells were cultured on gelatin-coated plates in ESC media Knockout D-MEM (Invitrogen, Waltham, MA) with 15% FBS, 0.1 mM MEMnon-essential amino acids, 2 mM GlutaMAX, 0.1 mM 2-mercaptoethanol (Sigma) and 1000 units/ml of ESGRO (Chem-icon). ES cells are fed daily and passaged every two days by trypsinization. Mitotic cells were synchronized by adding 100 ng/mL of Nocodazole for 6 hr followed by shake-off. For endogenously-tagged mAID-TBP cells, TBP degradation was performed by addition of indole-3-acetic acid (IAA) at 500 μM final concentration for 6 hours.

### Cas9-mediated endogenous knock-ins

Cas9-mediated endogenous knock-ins were performed as previously described (11). Briefly, mouse ES cells were transfected with 0.5 μg of the Cas9 vector (containing the specific guide RNA and the Venus coding sequence) and with 1 μg of the donor repair vector using Lipofectamine 3000. The transfected cells were sorted for Venus expressing cells the next day. Sorted cells were plated and grown on gelatin-coated plates under dilute conditions to obtain individual clones. After one week, individual clones were isolated and were used for direct cell lysis PCR using Viagen DirectPCR solution. PCR positive clones were further grown on gelatin-coated plates and cell lysates were obtained for Western blot analysis. Homozygous and heterozygous Halo-tagged TBP cell lines were generated, one homozygous of which was further tested by teratoma assay and was performed by Applied Stem Cell. We also generated homozygous mAID-TBP cell lines, one of which (C94) was further used to endogenously knock-in HaloTag to the Rpb1 locus, the largest subunit of RNA Polymerase II (C64 cell line). All primers and guide RNAs used to generate knock-ins are listed in Supplemental Table 1.

### Live cell and fixed sample imaging

Live-cell imaging experiments including FRAP were performed on cells grown on gelatin-coated glass bottom microwell dishes (MatTek #P35G-1.5–14-C) and labeled with the Halo-ligand dye JF549 at 100 nM concentration for 30 minutes. To remove unbound ligand, cells were washed 3x with fresh media for 5 minutes each. Epi-fluorescence time lapse imaging was performed on Nikon Biostation IM-Q equipped with a 40x/0.8 NA objective, temperature, humidity and CO2 control, and an external mercury illuminator. Images were collected every 2 minutes for 12 hr. Confocal live-cell imaging was performed using a Zeiss LSM 710 confocal microscope equipped with temperature and CO2 control. For fixed imaging, labeled cells were fixed with 4% PFA for 10 minutes in room temperature and were washed with 1x PBS prior to imaging. Standard immunofluorescence was performed on labeled cells using 4% PFA. Quantification of chromosome enrichment was performed using Fiji.

### FRAP

FRAP was performed on Zeiss LSM 710 confocal microscope with a 40x/1.3 NA oil-immersion objective and a 561 nm laser as previously described (11). Bleaching was performed using 100% laser power and images were collected at 1 Hz for the indicated time. FRAP data analysis was performed as previously described (26). For each cell line, we collected 10 cells for technical replicates in one experiment, which was repeated for a total of three biological replicates (30 cells total).

### Single molecule imaging – Slow tracking

Cells were labeled with JF549 at 10 pM for 30 minutes, and washed 3x with fresh media for 5 minutes each to remove unbound ligand. Cells were imaged in ESC media without phenol-red. A total of 600 frames were collected for imaging experiments at 500 ms frame rate (2 Hz) and were repeated 3x for biological replicates, with each experiment consisting of 10 cells for technical replicates each. Data are represented as mean over experimental replicates (30 cells total) ± SEM. Imaging experiments were conducted as described previously (11) on a custom-built Nikon TI microscope equipped with a 100x/NA 1.49 oil-immersion TIRF objective (Nikon apochromat CFI Apo SR TIRF 100x Oil), EM-CCD camera (Andor iXon Ultra 897), a perfect focusing system (Nikon) and a motorized mirror to achieve HiLo-illumination (47). Bound molecules were identified using SLIMfast as previously described (11). The length of each bound trajectory, corresponding to the time before unbinding or photobleaching, was determined and used to generate a survival curve (fraction still bound) as a function of time. A two-exponential function was then fitted to the survival curve, and the residence time was determined as previously described (11). The apparent half-life in seconds of bound Halo-Pol II was calculated by dividing the natural log of 2 with the photobleaching-corrected *k*_*off*_ (ln2/*k*_*off*_) extracted from the two-exponential function fitting. Photobleaching correction is performed by subtracting the apparent *k*_*off*_ of H2B-Halo from the apparent *k*_*off*_ of Pol II samples as previously described (11, 26).

### Single molecule imaging – Fast tracking

Cells were labeled with photo-activatable PA-JF549 at 25 nM for 30 minutes and washed 3x with fresh media for 5 minutes each to remove unbound ligand. Cells were imaged in ESC media without phenol-red. A total of 20,000 frames were collected for at 7.5 ms frame rate (133 Hz) and were repeated 3x for biological replicates, with each experiment consisting of eight cells of technical replicates. Data are represented as mean over experimental replicates (24 cells total) ± standard error of means. Single molecules were localized and tracked using SLIMfast, a custom-written MATLAB implementation of the MTT algorithm (48), using the following algorithm settings: Localization error: 10-6.25; deflation loops: 0; Blinking (frames); 1; maximum number of competitors: 3; maximal expected diffusion constant (µm2/s): 20. The fraction of bound molecules was determined as previously described using the following fit paramerters: TimeGap, 7.477; Gaps Allowed, 1; Jumps to consider, 4; Model fit type, CDF; Localization error, 0.045 (11, 26, 27) using Spot-On (source code freely available at https://gitlab.com/tjian-darzacq-lab/spot-on-matlab). The general statistics for image analysis (number of detections, number of total trajectories, and number of trajectories ≥3) for all spaSPT data are listed in Supplemental Table 2.

### ChIP-seq

ChIP-seq was performed as described previously (49). Briefly, mouse ES cells (JM8.N4) were cross-linked using 1% formaldehyde for 5 minutes at room temperature and washed with PBS before quenching with 125mM Glycine in PBS for 5 minutes. Cells were scraped, collected by centrifugation, and resuspended in 200 μL of Lysis buffer (1% SDS, 10mM EDTA, 50 mM Tris-HCl (pH 8.1), protease inhibitors) for 10 minutes on ice. Lysates were diluted 10-fold in cold ChIP dilution buffer (1% Triton X-100, 20 mM Tris-HCl (pH 8.1), 2mM EDTA, 150 mM NaCl, 5 mM CaCl_2_). Mnase (5 μL of 0.2 U/μL stock) was added to the lysate and samples were incubated at 37C for 15 minutes. Digested was quenched by adding EDTA (final concentration of 10 mM) and EGTA (final concentration 20 mM). Samples were sonicated using Branson digital sonifier at 40 seconds on time at 30% power with 2.5 seconds on and 5 seconds off cycles. Lysates were cleared by centrifugation and used as input for ChIP by adding *α*-TBP antibody (Abcam #ab51841) and incubating overnight. Protein-G magnetic beads were used to immuno-capture bound fragments, which were then washed as described previously (49), and DNA was purified by ethanol extraction. Two replicates were performed, and each sample and replicate was sequenced using one lane of Illumina Hi-Seq 2500 for 50 bp paired-end reads. Reads were mapped on mm10 genome build using Bowtie2 with the following parameters: --no-unal --local --very-sensitive-local --no-discordant --no-mixed --contain --overlap --dovetail -- phred33–I 10 –X 2000. Downstream analyses were performed using DeepTools suite (50). Data is deposited in GEO under the accession number GSE109964.

### Chr-RNA-seq

Chromatin associated nascent RNA was extracted as previously described (51). Briefly, 10 million celsl were collected for each sample as described in Figure 6C. Each sample was washed with ice-cold PBS and lysed with 800 μL of Buffer A (10 mM HEPES, pH 7.9, 10 mM KCl, 1.5 mM MgCl_2_, 0.34 M Sucrose, 10% Glycerol, 1 mM DTT, 0.1% Triton X-100, protease inhibitors) as described. Nuclei were subjected to consecutive biochemical fractionation with incubations for 15 minutes each with centrifugation and collection in between each fractionation. Nuclei were incubated 3 times for 15 minutes with Buffer B (9 mM EDTA, 20 mM EGTA, 1 mM DTT, 0.1% Triton X-100, protease inhibitors), then 2 times for 15 minutes each with Buffer B+ (20 mM EDTA, 20 mM EGTA, 2mM spermine, 5 mM spermidine, 1 mM DTT, 0.1% Triton X-100, protease inhibitors). Following fractionation, RNA from the insoluble chromatin was extracted with Trizol and prepared for sequencing as previously described (51). Sequencing was performed on one lane of Hi-Seq 4000 with 50 bp single-end reads.

### Chr-RNA-seq analysis

The sequenced reads were mapped to either the mm10 genome build or ERCC92 build for the spike-in controls using Tophat with the following parameters: library-type = fr-firststrand, b2-very-sensitive, no-coverage-search. The mapped reads were de-duplicated using the samtools rmdup function and then scaled using the scaling index calculated from the total number of reads that mapped to the ERCC92 build. The scaling equation is 1000/# of de-duplicated ERCC-mapped reads (Supplemental Table 3). Bigwig files were generated using DeepTools suite bamCoverage function, and reads were extended for 200 bp. Intronic transcripts per million (TPM) counts for each gene were calculated using Bioconductor R, which were then inputted for principal component analysis. Scatter plots were generated using R ggplots package. TSS plots, heatmaps, and k-means clustering analyses were generated with combined replicates using DeepTools suite (50). Data is deposited in GEO under the accession number GSE109964.

